# Serum Bile Acid Dysregulation in Polycystic Ovary Syndrome: Quantitative Insights from Mass Spectrometry-Based Profiling

**DOI:** 10.64898/2026.02.27.708465

**Authors:** Jalpa Patel, Hiral Chaudhary, Sonal Panchal, Bhavin Parekh, Rushikesh Joshi

## Abstract

**Background:** Polycystic ovary syndrome (PCOS) is a prevalent endocrine disorder with substantial metabolic comorbidities, including obesity, insulin resistance, and dyslipidaemia. Beyond their classical digestive role, bile acids (BAs) function as metabolic signalling molecules that regulate glucose and lipid homeostasis and inflammation through receptors such as the farnesoid X receptor (FXR) and Takeda G-protein receptor 5 (TGR5). However, bile acid dysregulation in PCOS remains inadequately characterised.

**Methods:** Targeted serum bile acid profiling was performed in PCOS (n = 86) and healthy controls (n = 60) using a validated LC-MS/MS method. Individual bile acids were quantified and classified into primary, secondary, and conjugated forms. Multivariate analyses were applied to identify group-level metabolic patterns. Functional bile acid indices reflecting hepatic conjugation and microbial transformation were calculated. Correlation analyses assessed between bile acids and clinical variables.

**Results:** PCOS women exhibited significantly higher serum levels of cholic acid and conjugated bile acids. Multivariate analyses revealed distinct bile acid signatures differentiating PCOS from controls, with deoxycholic acid, taurocholic acid, and cholic acid contributing most strongly to group separation. Pathway-based indices demonstrated an expanded conjugated bile acid pool, an increased conjugated-to-unconjugated bile acid ratio, and altered secondary-to-primary bile acid balance in PCOS. Several bile acids showed significant associations with androgen levels and gonadotropin ratios.

**Conclusion:** PCOS is characterised by coordinated alterations in bile acid metabolism, including hepatic synthesis, conjugation, and gut microbial transformation, highlighting bile acids as integrative metabolic signals linking endocrine and metabolic dysfunction in PCOS.

## Introduction

Polycystic ovary syndrome (PCOS) is a prevalent and complex hormonal disorder that affects up to 15% of women of reproductive age (1)(2). It presents with clinical and biochemical signs such as hyperandrogenism, ovulatory problems, and polycystic ovaries (3). Beyond its impact on fertility, PCOS is increasingly recognised as a systemic metabolic condition, often linked to insulin resistance, dyslipidaemia, abdominal obesity, and elevated risks of type 2 diabetes, cardiovascular disease, and non-alcoholic fatty liver disease (NAFLD)(4)(5)(6). Despite its widespread occurrence and complex causes, the exact molecular pathways linking the metabolic and reproductive aspects of PCOS remain incompletely understood.

Bile acids (BAs), once mainly considered metabolic by-products of cholesterol breakdown, crucial for lipid absorption, are now recognised as key signalling molecules that influence metabolic and hormonal pathways. Produced in the liver and subsequently modified by the gut microbiota, bile acids exert widespread systemic effects by activating nuclear receptors like the farnesoid X receptor (FXR) and membrane receptors such as Takeda G-protein receptor 5 (TGR5) (7)(8). These intricate signalling pathways play a critical role in regulating physiological processes, including glucose control, lipid metabolism, energy expenditure, and inflammation— many of which are often impaired in PCOS.

Recent research indicates that alterations in bile acid metabolism may both reflect and influence the metabolic disorders associated with PCOS (9). Disruption of the gut-liver axis and enterohepatic bile acid circulation has been strongly associated with insulin resistance, fatty liver disease, and endocrine issues—hallmarks of PCOS (10). Nevertheless, studies on bile acid profiles in women with PCOS are limited. Only a few have investigated circulating bile acid levels, and their findings are inconsistent. These differences likely stem from variations in testing techniques, study populations, and other clinical factors.

Mass spectrometry techniques, particularly liquid chromatography-tandem mass spectrometry (LC-MS/MS), provide remarkable sensitivity and specificity for detecting individual bile acid types, including primary, secondary, conjugated, and unconjugated forms (11)(12). This targeted method allows for a thorough and accurate evaluation of bile acid metabolism, surpassing traditional biochemical assays and untargeted metabolomics approaches. In our study, we used targeted LC-MS/MS to quantify serum bile acids in women with PCOS and healthy controls. By providing precise quantitative data on circulating bile acids, this research aims to enhance current knowledge and investigate the potential of bile acids as metabolic biomarkers or contributors to PCOS pathophysiology.

## Methods

### Participants Recruitment and Ethical Approval

We recruited 146 women aged 17 to 40 for this study. Among them, 86 were diagnosed with PCOS based on the Rotterdam Criteria, which conforms to the ESHRE/ASRM consensus. This diagnosis required at least two of the following three criteria: clinical or lab evidence of androgen excess, chronic anovulation, or polycystic ovaries seen via transvaginal ultrasonography. The control group consisted of 60 women without endocrine disorders and not using hormonal contraceptives. All women with PCOS were selected from patients at Dr. Nagori’s Institute for Infertility and IVF in Ahmedabad. The Ethical Committee of Gujarat University (GU-IEC(NIV)/02/Ph.D./007) approved the study to ensure compliance with ethical standards.

### Sample Collection and Biochemical Analyses

Blood specimens were collected from participants during the follicular phase (days 2-5) of their menstrual cycle after an overnight fast. This timing was intentional, designed to enhance conditions for biochemical evaluations and serum metabolomic analysis. Serum samples for the research were collected between January 2023 and December 2024, representing metabolic profiles from both PCOS and healthy controls, and participants were informed of the study’s purpose. After collection, the specimens were centrifuged at 2500 rpm for 15 minutes, and the serum was stored at −80°C until analysis. Biochemical investigations included measuring luteinizing hormone (LH), follicle-stimulating hormone (FSH), and 17β-estradiol, along with thyroid-stimulating hormone (TSH) and prolactin (PRL) to rule out thyroid and prolactin disorders. Testosterone (T) and dehydroepiandrosterone sulfate (DHEA-S) levels were also measured to evaluate hyperandrogenemia, using chemiluminescent assays at an accredited clinical laboratory.

### Chemicals and Materials

Cholic acid (CA), Chenodeoxycholic acid (CDCA), Deoxycholic acid (DCA), Lithocholic acid (LCA), Taurocholic acid (TCA), Glycocholic acid (GCA), and Taurodeoxycholate (TDCA) were purchased from Sigma-Aldrich (St. Louis, MO, USA). Cholic acid-d4, as an internal standard, was purchased from Cayman Chemicals (Michigan, USA). LC/MS-grade water, ammonium acetate, and methanol were purchased from J.T. Baker.

### Sample Preparation

Initially, 50 microliters of serum were combined with 300 μL of methanol and 20 μL of internal standard solution in centrifuge tubes. After thorough mixing, the samples were cooled at −20°C for 20 minutes to promote protein precipitation. Subsequently, centrifugation was performed at 13,000 g for 12 minutes. Each sample was transferred into a glass LC-MS vial and dried using a vacuum evaporator. The resulting dry residues were reconstituted with 300 μL of a methanol-water mixture (60:40). Finally, the samples were filtered through a 0.22 μm nylon filter before 10 μL of the supernatant was injected into the LC-MS/MS system.

### Chromatography conditions LC conditions

The analysis was performed on a Bruker Elute UHPLC system controlled by Hystar 5.0 SR1 software. Mobile phase A consisted of 250 mM ammonium acetate (pH 5.3) in a 1:49 ratio with water, while mobile phase B was 250 mM ammonium acetate (pH 5.3) in a 1:49 ratio with methanol. Various bile acid species were separated on a Phenomenex Kinetex C18 HPLC column (150 x 4.6 mm, 2.6 μm) using a 12-minute LC program. The LC conditions were set to maintain 80% mobile phase B for 0.5 minutes, followed by a linear increase from 80% to 98% mobile phase B over 7 minutes. Subsequently, the system was held at 98% mobile phase B for 2.4 minutes before equilibrating with 80% mobile phase B for the remaining time. The flow rate was 0.4 mL/min, and the column temperature was maintained at 30°C.

### MS / MS conditions

Quantification was performed in multiple reaction monitoring (MRM) mode with electrospray negative ionization on the Bruker Daltonics AmaZon Speed™ ion trap mass spectrometer. Source- and compound-specific parameters were optimized to achieve the highest signal response with direct infusion over a scan range of 100-1000 m/z. The source-specific parameters were as follows: nebulizer, 29.0 psi; dry temperature, 126.9 °C; dry gas, 10.0 L/min; capillary voltage, 4500 V. Data were collected and analysed using Data Analysis 5.2, provided by Bruker Daltonics Gmbh. A calibration curve was constructed by plotting the peak area ratios of each analyte to the isotope-labelled internal standard against the amount of analyte.

### Validation Parameters

Method validation includes assessing linearity, LLOD, LLOQ, apparent recovery, and matrix effect. To evaluate linearity, a calibration curve was created by plotting the peak area ratios of each bile acid to the isotope-labelled internal standard against the analyte concentrations. A series of dilutions for each bile acid generated a 6-point calibration curve with concentrations of 5 μg/ml, 10 μg/ml, 15 μg/ml, 20 μg/ml, 25 μg/ml, and 30 μg/ml. The internal standard, cholic acid d4, was kept at a constant 15 μg/ml for calculating the area ratios.

### Apparent recovery and matrix effect

To evaluate the apparent recovery and matrix effect, synthetic serum samples were spiked with bile acids at concentrations of 5μg/ml, 15μg/ml, and 30μg/ml (n = 3 for each level). Unextracted samples were prepared with methanol, while post-extracted samples involved spiking bile acids into the extracted synthetic serum. Pre-extracted samples were created by adding bile acids to the synthetic serum before the extraction process. Apparent recovery was calculated by comparing pre-extracted to post-extracted samples, and the matrix effect was assessed by comparing post-extracted samples with unextracted ones.

**Apparent recovery %**= Peak area ratio of [ pre-extracted sample/post-extracted of sample] x 100

**Matrix effects %**= Peak area ratio of [ post-extracted sample/ Unextracted of sample] x 100

### Statistical analysis

The statistical analysis was conducted using SPSS version 20.0 (SPSS Inc., Chicago, IL, USA) and GraphPad Prism 8.0 (GraphPad Software, Inc., San Diego, CA). The distribution of each continuous variable was evaluated with the Kolmogorov–Smirnov test. Data were expressed as mean ± SD or median (interquartile range, 25th–75th percentile), depending on the normality of distribution. Differences between the two groups were assessed using Student’s t-test and the Mann-Whitney U test. Spearman’s rank correlation was used for correlation analysis. To compare patients with PCOS to controls in multivariate analyses, MetaboAnalyst 6.0 was employed. Principal component analysis (PCA) and orthogonal partial least squares-discriminant analysis (OPLS-DA) were performed on the metabolomics data. The variable importance in projection (VIP) scores from the first principal component in OPLS-DA were calculated. Metabolites with VIP > 1.0 and P-value < 0.05 were considered significantly different in univariate analysis. All statistical tests were two-sided, with P < 0.05 considered statistically significant.

## RESULTS

### Clinical and Biochemical Characteristics of the Study Cohort

The targeted serum bile acid analysis revealed significant biochemical and clinical differences between women with PCOS (n=86) and control subjects (n=60) (Table 1). Women with PCOS were older than controls (28.72 ± 4.14 years vs. 25.03 ± 5.16 years, p < 0.0001) and had a higher BMI (26.22 ± 5.00 kg/m^2^ vs. 23.58 ± 4.78 kg/m^2^, p = 0.001).

**Table 1.** Demographic and biochemical parameters of the study participants.

| Demographic and Biochemical parameters | PCOS (n=86) | Control (n=60) | p-value |
| --- | --- | --- | --- |
| Age (year) | $28.72 \pm 4.14$ | $25.03 \pm 5.16$ | $<0.0001$ |
| BMI (kg/m<sup>2</sup>) | $26.22 \pm 5.00$ | $23.58 \pm 4.78$ | 0.001 |
| Testosterone (ng/dl) | $30.41 \pm 21.54$ | $17.31 \pm 8.15$ | $<0.0001$ |
| Estradiol (pg/ml) | $73.28 \pm 47.83$ | $68.97 \pm 27.80$ | 0.53 |
| LH (mIU/ml) | $7.10 \pm 4.25$ | $5.35 \pm 4.45$ | 0.019 |
| FSH (mIU/ml) | $8.66 \pm 6.65$ | $8.23 \pm 6.21$ | 0.69 |
| PRL (ng/ml) | $15.85 \pm 12.42$ | $13.77 \pm 8.67$ | 0.26 |
| LH/FSH ratio | $1.05 \pm 0.90$ | $0.97 \pm 1.14$ | 0.63 |
| DHEA-S (ug/dl) | $140.86 \pm 73.31$ | $149.96 \pm 60.47$ | 0.43 |
| TSH (uIU/ml) | $1.85 \pm 0.96$ | $2.03 \pm 1.20$ | 0.32 |
| Fasting Glucose | $83.92 \pm 15.44$ | -- | -- |
| Postprandial Glucose | $111.94 \pm 29.11$ | -- | -- |
| Fasting Insulin | $9.87 \pm 3.60$ | -- | -- |
| Postprandial Insulin | $86.59 \pm 69.99$ | -- | -- |
| Homa IR | $2.11 \pm 0.78$ | -- | -- |

Testosterone levels were significantly elevated in the PCOS group compared to controls (30.41 ± 21.54 ng/dl vs. 17.31 ± 8.15 ng/dl, p < 0.0001). Estradiol levels were similar between the groups (73.28 ± 47.83 pg/ml in PCOS vs. 68.97 ± 27.80 pg/ml in controls; p = 0.53). LH levels were higher in the PCOS group (7.10 ± 4.25 mIU/ml) compared to controls (5.35 ± 4.45 mIU/ml, p = 0.019), while FSH levels did not differ significantly (8.66 ± 6.65 mIU/ml for PCOS vs. 8.23 ± 6.21 mIU/ml for controls, p = 0.69). The LH/FSH ratio was also comparable between groups (1.05 ± 0.90 for PCOS vs. 0.97 ± 1.14 for controls, p = 0.63). PRL levels (15.85 ± 12.42 ng/ml for PCOS vs. 13.77 ± 8.67 ng/ml for controls, p = 0.26), DHEA-S levels (140.86 ± 73.31 μg/dl for PCOS vs. 149.96 ± 60.47 μg/dl for controls, p = 0.43), and TSH levels (1.85 ± 0.96 μIU/ml for PCOS vs. 2.03 ± 1.20 μIU/ml for controls, p = 0.32) were similar between the groups.

For the PCOS group, fasting glucose was 83.92 ± 15.44 mg/dl, postprandial glucose was 111.94 ± 29.11 mg/dl, fasting insulin was 9.87 ± 3.60 μIU/ml, postprandial insulin was 86.59 ± 69.99 μIU/ml, and HOMA-IR was 2.11 ± 0.78. These glucose and insulin measures were not available for the control group.

### LC-MS/MS Optimization and Chromatographic Validation of Bile Acids

The proposed LC-MS/MS method enables quantification of seven bile acid species and their metabolites, including taurine and glycine conjugates, from 50 μL of serum in a 12-minute run. The key fragment ions used for quantification were carefully chosen: m/z 74, representing [NH2 CH2 COO] −, for glycine-conjugated bile acids, and m/z 80, representing [SO3] −, for taurine-conjugated bile acids. Since specific fragment ions are lacking for unconjugated bile acids, their quantification was achieved by monitoring the same m/z for both the parent and fragment ions, following published methods (13).

The assay showed linearity for all tested bile acid species within a concentration range of 5 μg/mL to 30 μg/mL, with R^2^ values between 0.991 and 0.998. The LLOD and LLOQ for each bile acid were both set at 5 μg/mL. No carryover was detected. A six-point calibration curve was created for each bile acid (see Supplementary Figures 1 to 7) and used to quantify serum samples.

Apparent recovery was evaluated at three different levels (n = 3 per level) with synthetic serum. The recoveries ranged between 87.8% and 107.0% (Table 2). The matrix effect was also evaluated at three different levels (n = 3 per level) (Table 3). Ion enhancement was seen with CDCA and DCA at low concentrations; ion suppression was seen with LCA and GCA at high concentrations (Table 3).

**Table 2.** The Result from apparent recovery study with synthetic serum.

| Bile acids | Apparent recovery % |  |  |
| --- | --- | --- | --- |
|  | 5µg/ml | 15µg/ml | 30µg/ml |
| Cholic acid | 104.6 | 96.7 | 100.4 |
| Chenodeoxycholic acid | 98.6 | 102.7 | 95.4 |
| Deoxycholic acid | 100.1 | 95.0 | 99.7 |
| Lithocholic acid | 98.2 | 93.7 | 89.6 |
| Taurocholic acid | 103.6 | 94.9 | 90.6 |
| Glycocholic acid | 87.8 | 98.4 | 101.0 |
| Taurodeoxycholate | 90.8 | 106.7 | 107.0 |

**Table 3.**
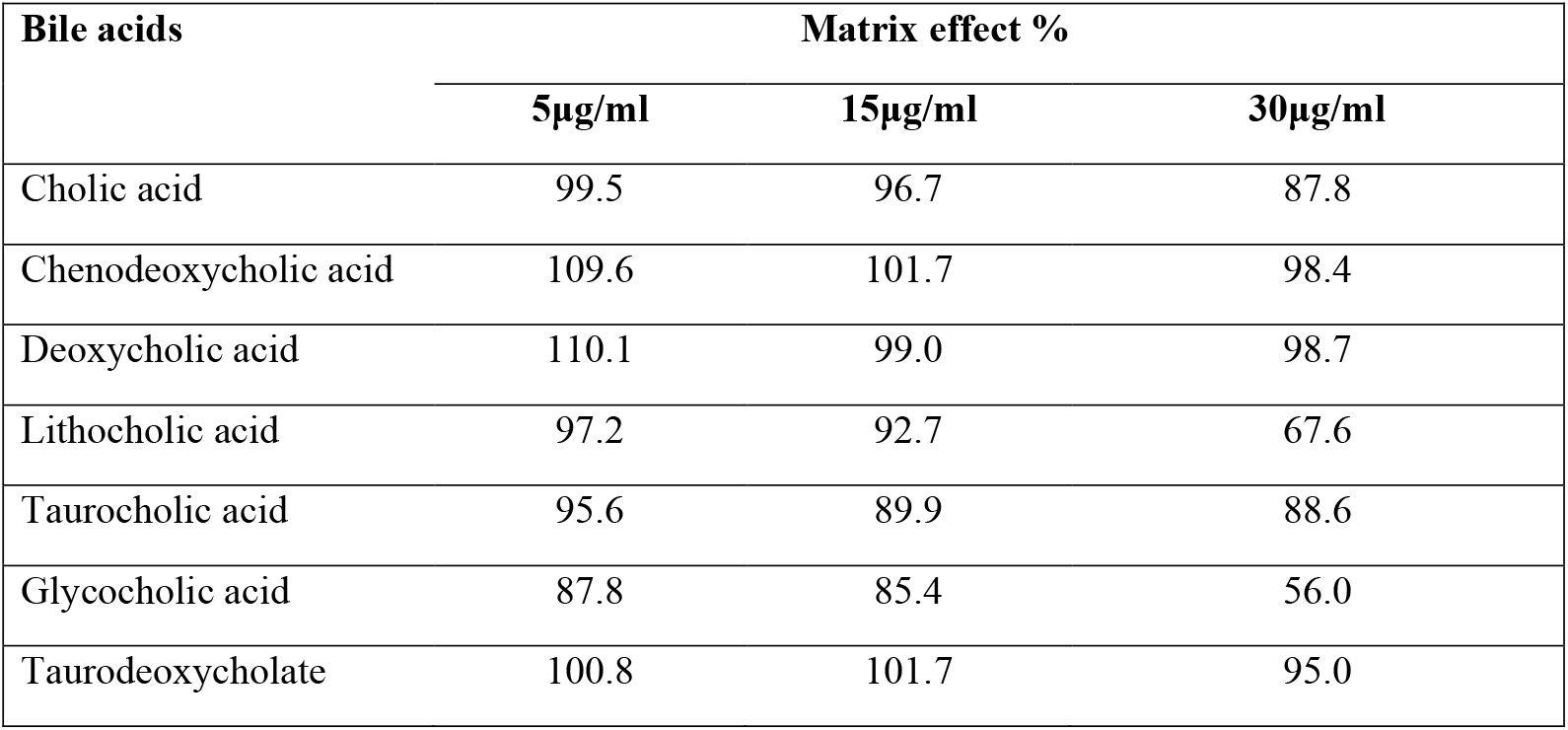
Result from matrix study with synthetic serum.

### Distribution of Individual Serum Bile Acids Across Study Groups

As we have developed a method for quantifying serum bile acids, we have prepared a calibration curve in ug/ml. After getting the regression equation for each bile acids to quantify unknown amount of bile acid from serum sample, we have converted ug/ml into the suitable unit of bile acid measurement that is nmol/ml with the use of formular: Concentration in nmol/ml= (concentration in μg/ml x 1000/ Molecular weight of bile acid (g/mol).

This study compared serum bile acid levels between women with PCOS (n=86) and a control group (n=60) (Table 4). The results showed significant differences in several bile acids. Among primary bile acids, cholic acid (CA) was notably higher in the PCOS group (0.87 ± 0.82 nmol/ml) than in the control group (0.56 ± 0.55 nmol/ml; p=0.01). However, chenodeoxycholic acid (CDCA) levels did not differ significantly between the two groups, with values of 5.35 ± 4.88 nmol/ml in the PCOS group and 4.77 ± 4.13 nmol/ml in controls (p=0.45).

**Table 4.** Comparison of Serum Bile acid level in the PCOS with the Control group.

| Serum Bile acids | m/z | PCOS (n=86) | Control (n=60) | p-value |
| --- | --- | --- | --- | --- |
| **Primary Bile Acids** |  |  |  |  |
| Cholic acid (CA) nmol/ml | 407.33<sup>-1</sup> | 0.87 ± 0.82 | 0.56 ± 0.55 | **0.01** |
| Chenodeoxycholic acid (CDCA) nmol/ml | 391.35<sup>-1</sup> | 5.35 ± 4.88 | 4.77 ± 4.13 | 0.45 |
| **Secondary Bile Acids** |  |  |  |  |
| Deoxycholic acid (DCA) nmol/ml | 391.32<sup>-1</sup> | 1.91 ± 2.57 | 1.44 ± 2.26 | 0.25 |
| Lithocholic acid (LCA) nmol/ml | 375.34<sup>-1</sup> | 20.76 ± 16.50 | 18.41 ± 16.78 | 0.40 |
| **Conjugated Bile acids** |  |  |  |  |
| Taurocholic acid (TCA) nmol/ml | 514.33<sup>-1</sup> | 7.13 ± 2.17 | 5.39 ± 0.35 | **<0.0001** |
| Glycocholic acid (GCA) nmol/ml | 464.36<sup>-1</sup> | 6.31 ± 2.35 | 5.19 ± 0.27 | **<0.001** |
| Taurodeoxycholate (TDCA) | 498.31<sup>-1</sup> | 6.58 ± 8.99 | 3.81 ± 6.86 | **0.04** |

For secondary bile acids, DCA and LCA levels were similar across groups. DCA was 1.91 ± 2.57 nmol/ml in the PCOS group versus 1.44 ± 2.26 nmol/ml in controls (p=0.25), and LCA was 20.76 ± 16.50 nmol/ml versus 18.41 ± 16.78 nmol/ml (p=0.40). Conjugated bile acids showed significant differences. TCA levels were higher in the PCOS group (7.13 ± 2.17 nmol/ml) than in controls (5.39 ± 0.35 nmol/ml), with p<0.0001. GCA was also elevated in PCOS women (6.31 ± 2.35 nmol/ml) compared to controls (5.19 ± 0.27 nmol/ml, p<0.001). Additionally, TDCA levels were increased in the PCOS group (6.58 ± 8.99 nmol/ml) versus controls (3.81 ± 6.86 nmol/ml; p=0.04).

### Multivariate Patterns Underlying Serum Bile Acid Variation in PCOS

To examine overall variation in serum bile acid profiles between the study groups, unsupervised and supervised multivariate analyses were performed (Figure 1). Principal component analysis (PCA) showed that the first two principal components accounted for a substantial proportion of the total variance (PC1: 45.6%, PC2: 27.0%) (Figure 1a). Samples from women with PCOS and control subjects exhibited partial overlap, indicating shared features of serum bile acid composition. However, a noticeable shift in the distribution and centroid positioning of the two groups suggested differences in the relative contribution of individual bile acids to overall variability. To further explore group-associated patterns, orthogonal partial least squares– discriminant analysis (OPLS-DA) was applied. The OPLS-DA score plot demonstrated separation primarily along the predictive component (T score [1], 14.4%), while the orthogonal component (Orthogonal T score [1], 30%) captured within-group variability unrelated to group classification (Figure 1b). The corresponding 95% confidence ellipses showed distinct clustering patterns for the PCOS and control groups, despite some overlap, reflecting heterogeneity in serum bile acid profiles. Variable importance in projection (VIP) analysis was subsequently used to identify bile acids contributing most strongly to the observed group-related variation (Figure 1c). Deoxycholic acid exhibited the highest VIP score, followed by taurocholic acid and cholic acid, with additional contributions from taurodeoxycholic acid and glycocholic acid. In contrast, chenodeoxycholic acid and lithocholic acid showed lower VIP values, indicating a relatively smaller contribution to group differentiation within the multivariate model.

**Figure 1.**
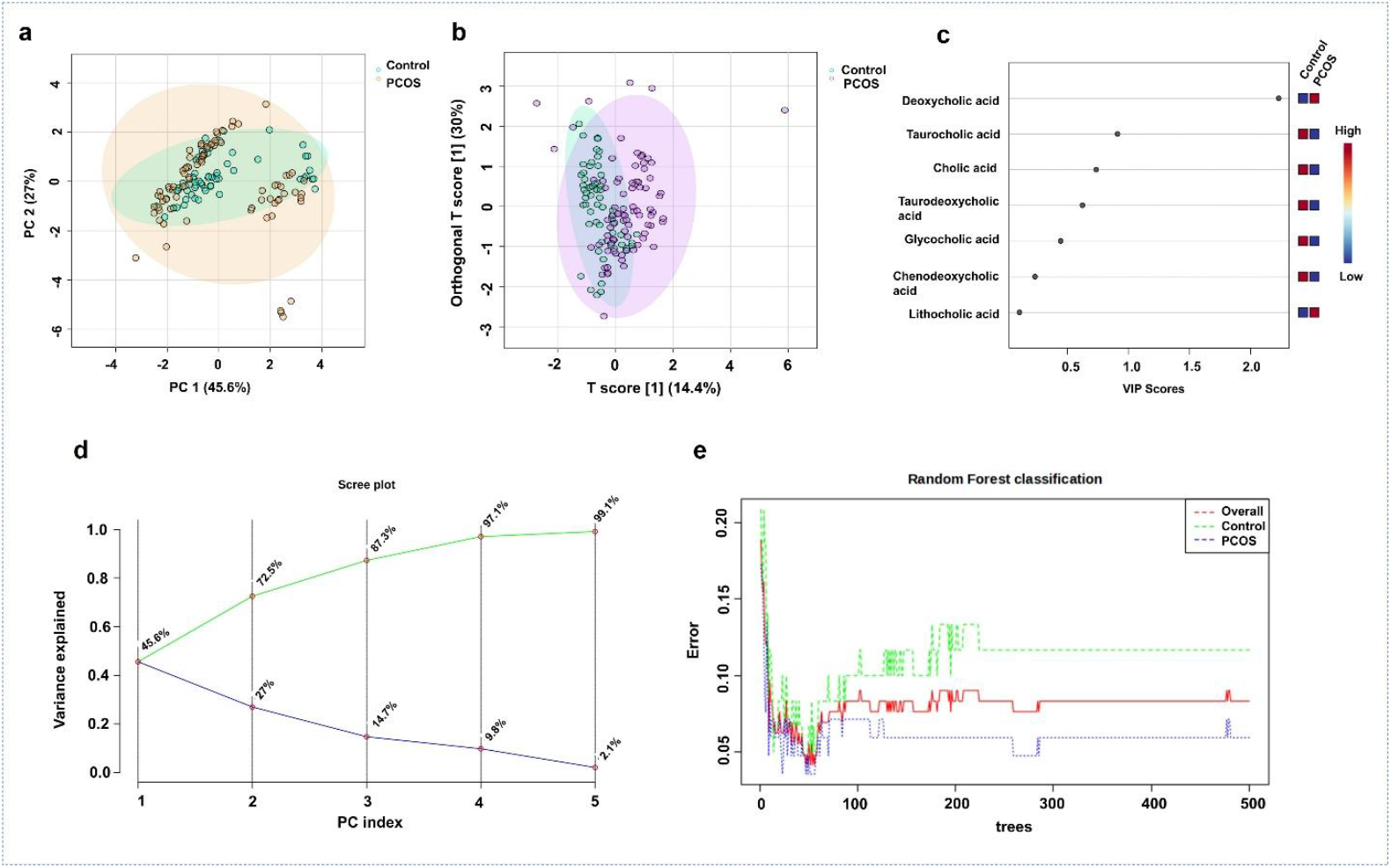
Multivariate characterization and validation of serum bile acid profiles in women with PCOS and control subjects. (a) PCA score plot shows serum bile acid profiles; first two components explain 45.6% (PC1) and 27.0% (PC2) of variance. Ellipses indicate 95% confidence for each group. (b) OPLS-DA score plot illustrates group differences, with 14.4% (T score [1]) and 30% (Orthogonal T score [1]) variance, and confidence ellipses. (c) VIP scores reveal key bile acids influencing group separation, with higher scores indicating greater impact. d) Scree plot shows the variance explained by principal components, with the first two capturing most of the total variance. (e) Random Forest error plot indicates stable, low error rates for PCOS and control groups across more decision trees, demonstrating the robustness and predictive power of serum bile acid profiles.

To further evaluate the robustness and dimensional structure of the multivariate models, additional validation analyses were performed. Examination of the scree plot revealed that the first principal component accounted for 45.6% of the total variance, with the second component contributing an additional 27.0%, for a total of over 70% of the variance in serum bile acid profiles (Figure 1(d)). Subsequent components contributed progressively smaller proportions of variance, indicating that the primary metabolic information was captured within a limited number of dimensions.

In parallel, Random Forest classification was used as an independent, non-parametric approach to assess the predictive stability of serum bile acid signatures. The classification error rapidly decreased and stabilised with increasing numbers of trees, with consistently lower error rates observed for the PCOS group compared to controls (Figure 1e). These findings provide independent confirmation of the discriminatory potential and robustness of bile acid–based metabolic signatures distinguishing PCOS from control subjects.

### Interrelationships among serum bile acids

Correlation analysis assessed the relationships among individual serum bile acids and their coordinated behavior within the circulating pool (Figure 2). Hierarchical clustering identified clear grouping patterns: primary bile acids and their conjugates clustered separately from secondary bile acids, indicating structured co-variation among subclasses. Strong positive correlations were found between cholic acid and its conjugates, glycocholic acid and taurocholic acid, reflecting their shared biosynthetic pathway and hepatic conjugation. Glycine- and taurine-conjugated bile acids also clustered closely, highlighting similarities in their biochemical properties and processing

**Figure 2.**
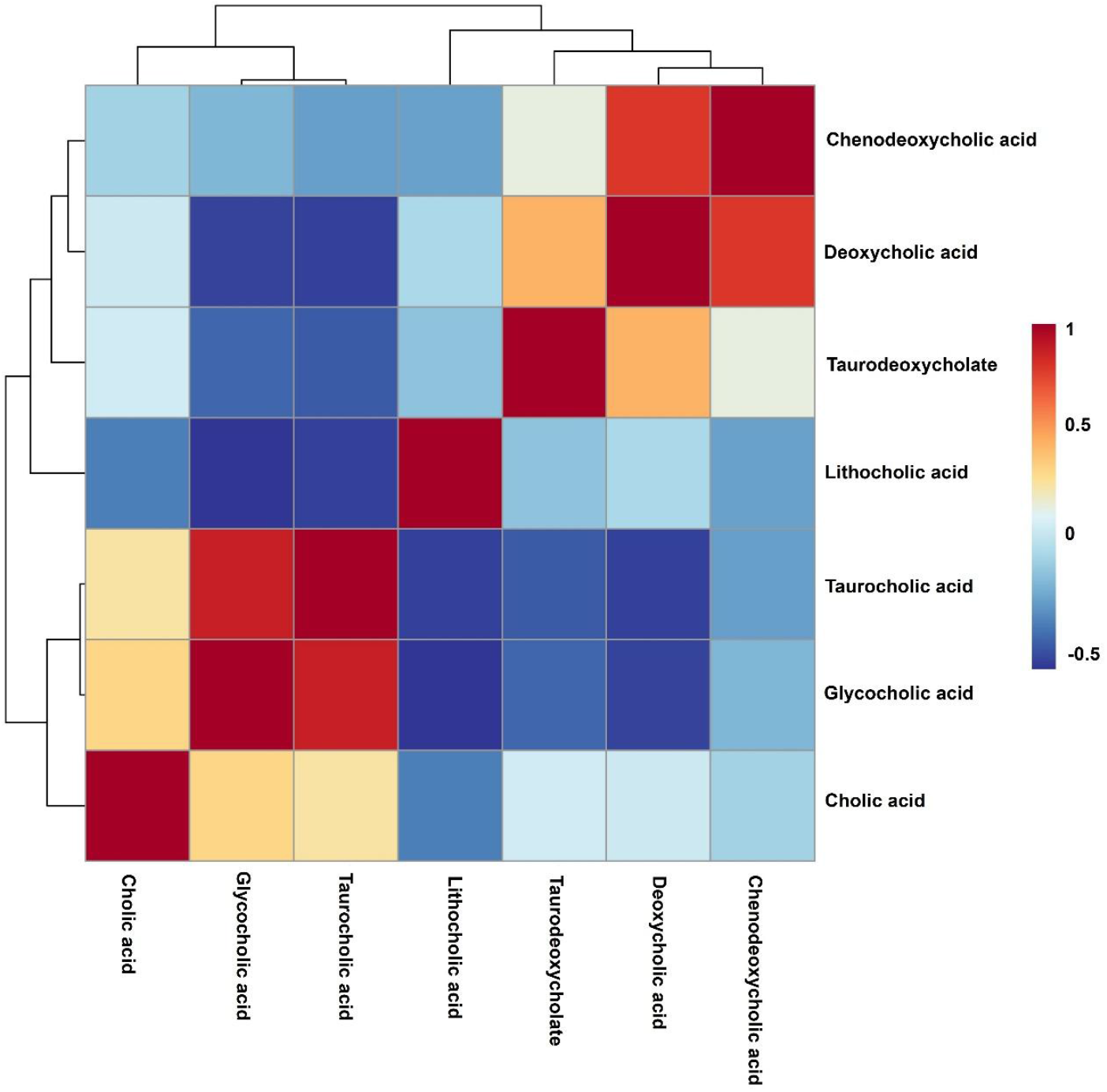
The correlation heatmap shows relationships among serum bile acids, with hierarchical clustering highlighting pairwise correlations. Color intensity indicates the correlation’s strength and direction (red, positive; blue, negative). Dendrograms reveal that primary and conjugated bile acids cluster separately from secondary acids.

In contrast, secondary bile acids, particularly deoxycholic acid and lithocholic acid, formed a separate cluster and showed inverse correlations with several conjugated bile acids. This pattern is consistent with the intestinal microbial transformation of primary bile acids into secondary bile acids. Taurodeoxycholate displayed an intermediate correlation pattern, clustering between conjugated and secondary bile acids, reflecting its position at the interface of hepatic conjugation and microbial modification. Overall, the correlation structure highlights coordinated variation in serum bile acids along the hepatic–intestinal axis and underscores the interconnectedness of bile acid biosynthesis, conjugation, and microbial conversion within the circulating bile acid profile.

### Serum Bile Acid Indices in Women with PCOS and Controls

To provide a functional overview of circulating bile acid metabolism, bile acids were grouped by hepatic origin and subsequent microbial transformation, and composite indices were derived (Figure 3). Primary bile acids, cholic acid (CA) and chenodeoxycholic acid (CDCA), are synthesised from cholesterol in the liver and undergo conjugation with glycine or taurine to form conjugated bile acids, including taurocholic acid (TCA), glycocholic acid (GCA), and taurodeoxycholic acid (TDCA). Following secretion into the intestinal lumen, conjugated bile acids may be deconjugated by gut microbial bile salt hydrolases, and primary bile acids can be further converted into secondary bile acids, such as deoxycholic acid (DCA) and lithocholic acid (LCA), through microbial 7α-dehydroxylation. A substantial proportion of bile acids is subsequently reabsorbed and returned to the liver via enterohepatic circulation (Figure 3(a)).

**Figure 3.**
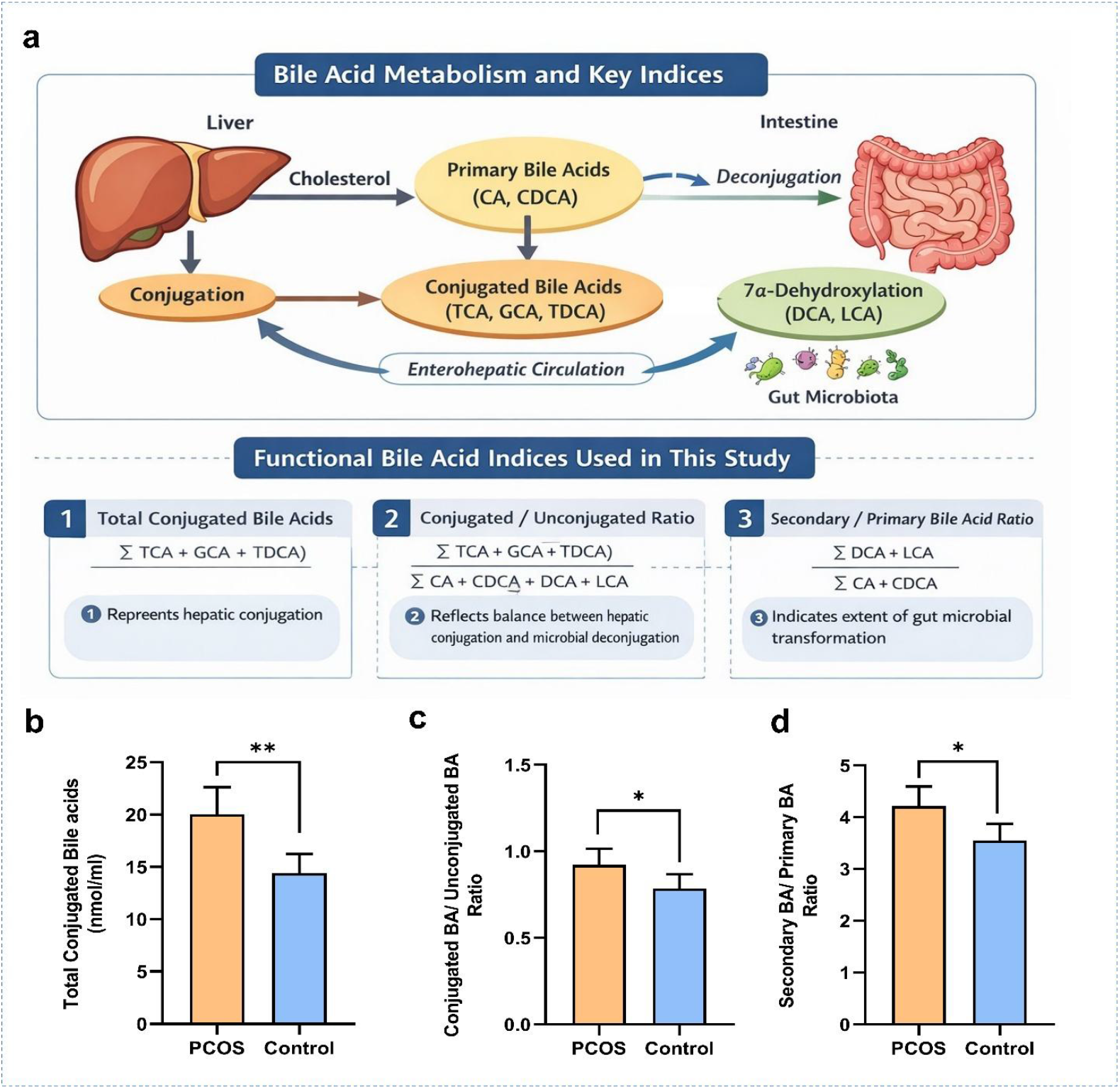
Conceptual overview and comparison of serum bile acid indices. (a) Schematic representation of hepatic bile acid synthesis, conjugation, gut microbial deconjugation and secondary bile acid formation, highlighting the bile acid pools and ratios used in this study. (b–d) Box-and-whisker plots comparing serum bile acid indices between PCOS and control groups, including total conjugated bile acids, conjugated-to-unconjugated bile acid ratio, and secondary-to-primary bile acid ratio.

Based on these metabolic pathways, three functional bile acid indices were calculated to summarise differences in serum bile acid composition between women with PCOS and control subjects. The total conjugated bile acid pool (Σ TCA + GCA + TDCA) was significantly higher in the PCOS group compared with controls (Figure 3(b)). The conjugated-to-unconjugated bile acid ratio, reflecting the relative contribution of conjugated bile acids within the circulating bile acid pool, was also increased in women with PCOS (Figure 3(c)). In addition, the secondary-to-primary bile acid ratio [(Σ DCA + LCA)/(Σ CA + CDCA)], representing the extent of conversion of primary bile acids into secondary bile acids, differed significantly between the PCOS and control groups (Figure 3(d)).

### Associations Between Serum Bile Acids and Clinical Parameters

The analysis of correlations between demographic and biochemical parameters and various bile acids in women with PCOS and control subjects revealed several interesting associations. Among primary bile acids, Cholic acid showed no significant correlations with age, BMI, testosterone, estradiol, TSH, LH, FSH, PRL, or the LH/FSH ratio in either group. However, in the PCOS group, a significant positive correlation with DHEAS was observed (r = 0.21, p = 0.05). Chenodeoxycholic acid showed a notable positive correlation with BMI in the control group (r = 0.28, p = 0.02) and with DHEAS in both the PCOS (r = 0.22, p = 0.03) and control groups (r = 0.32, p = 0.01). Furthermore, a significant negative correlation was observed between testosterone and Chenodeoxycholic acid in the control group (r = −0.26, p = 0.03) (Table 5).

**Table 5.**
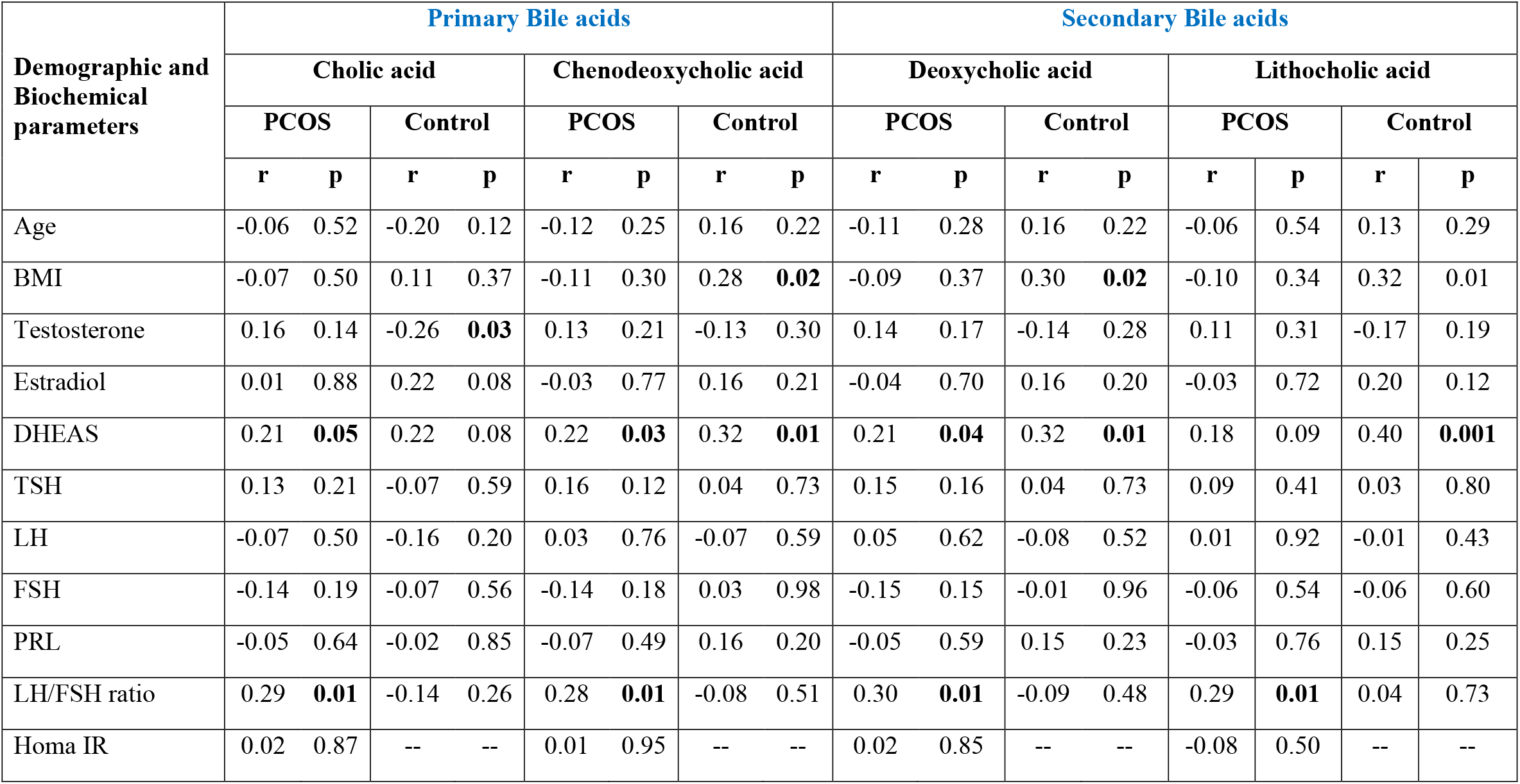
Correlation analysis between Bile acids (Primary and Secondary) and biochemical parameters.

The study examined the correlation between secondary bile acids and various biochemical parameters (Table 5). It was found that Deoxycholic acid showed significant positive correlations with BMI in the control group (r = 0.30, p = 0.02) and with DHEAS in both the PCOS (r = 0.21, p = 0.04) and control groups (r = 0.32, p = 0.01). Additionally, Deoxycholic acid exhibited a significant positive correlation with the LH/FSH ratio in the PCOS group (r = 0.30, p = 0.01). Lithocholic acid also displayed significant positive correlations with BMI in the control group (r = 0.32, p = 0.01) and with DHEAS in both the PCOS (r = 0.18, p = 0.09) and control groups (r = 0.40, p = 0.001). Furthermore, the LH/FSH ratio showed a significant positive correlation with Lithocholic acid in the PCOS group (r = 0.29, p = 0.01).

The study found no significant correlations between age, BMI, estradiol, TSH, LH, FSH, PRL, or the LH/FSH ratio and conjugated bile acids (taurocholic acid) in either group (Table 6). However, in the PCOS group, there was a significant positive correlation with testosterone (r = 0.28, p = 0.01). In the control group, BMI showed a significant negative correlation with taurocholic acid (r = −0.34, p = 0.01). Glycocholic acid showed a notable positive correlation with testosterone in the PCOS group (r = 0.21, p = 0.05) and with DHEAS in the control group (r = 0.27, p = 0.03). No significant correlations were observed with other parameters such as age, BMI, estradiol, TSH, LH, FSH, PRL, or the LH/FSH ratio in either group. Taurodeoxycholic acid did not exhibit any significant correlations with age, BMI, testosterone, estradiol, TSH, LH, FSH, or PRL in either the PCOS or control groups. However, a significant positive correlation was observed with the LH/FSH ratio in the PCOS group (r = 0.26, p = 0.01).

**Table 6.** Correlation analysis between Bile acids (Conjugated form) and biochemical parameters.

| Demographic and Biochemical parameters | Conjugated Bile acids |  |  |  |  |  |  |  |  |  |  |  |
| --- | --- | --- | --- | --- | --- | --- | --- | --- | --- | --- | --- | --- |
|  | Taurocholic acid |  |  |  | Glycocholic acid |  |  |  | Taurodeoxycholic acid |  |  |  |
|  | PCOS |  | Control |  | PCOS |  | Control |  | PCOS |  | Control |  |
|  | r | p | r | p | r | p | r | p | r | p | r | p |
| Age | 0.03 | 0.73 | -0.11 | 0.37 | -0.05 | 0.62 | 0.07 | 0.55 | -0.01 | 0.95 | -0.05 | 0.70 |
| BMI | -0.15 | 0.16 | -0.34 | **0.01** | -0.03 | 0.77 | 0.07 | 0.58 | -0.05 | 0.60 | 0.13 | 0.32 |
| Testosterone | 0.28 | **0.01** | 0.08 | 0.53 | 0.21 | 0.05 | -0.09 | 0.49 | 0.09 | 0.41 | -0.13 | 0.29 |
| Estradiol | 0.12 | 0.25 | -0.10 | 0.42 | -0.01 | 0.87 | 0.11 | 0.37 | 0.01 | 0.89 | 0.18 | 0.16 |
| DHEAS | 0.14 | 0.17 | -0.04 | 0.72 | 0.13 | 0.23 | 0.27 | **0.03** | 0.15 | 0.15 | 0.26 | **0.03** |
| TSH | -0.09 | 0.41 | -0.04 | 0.74 | 0.10 | 0.35 | 0.16 | 0.21 | 0.12 | 0.24 | 0.03 | 0.81 |
| LH | -0.05 | 0.60 | 0.13 | 0.32 | -0.08 | 0.44 | -0.12 | 0.35 | 0.04 | 0.67 | -0.10 | 0.41 |
| FSH | 0.01 | 0.32 | -0.01 | 0.91 | -0.12 | 0.27 | 0.01 | 0.94 | -0.12 | 0.27 | 0.05 | 0.67 |
| PRL | -0.01 | 0.98 | 0.05 | 0.65 | -0.05 | 0.63 | 0.14 | 0.28 | -0.09 | 0.37 | 0.17 | 0.17 |
| LH/FSH ratio | 0.10 | 0.32 | 0.04 | 0.73 | 0.26 | **0.01** | -0.17 | 0.18 | 0.18 | 0.10 | -0.14 | 0.26 |
| Homa IR | 0.02 | 0.83 | -- | -- | 0.01 | 0.88 | -- | -- | 0.02 | 0.81 | -- | -- |

## DISCUSSION

This study presents a comprehensive analysis of serum bile acid profiles in women with PCOS compared to healthy controls, revealing significant alterations in bile acid metabolism that may contribute to the pathophysiology of this complex endocrine disorder.

The findings show elevated conjugated bile acids, especially taurocholic acid, glycocholic acid, and taurodeoxycholate, as well as increased cholic acid, revealing new insights into PCOS-related metabolic issues. The key finding is the significant rise in conjugated bile acids in women with PCOS. Both conjugated primary bile acids, taurocholic acid and glycocholic acid, were higher in PCOS patients, consistent with previous studies (14)(15). Yang et al. also found increased glycocholic acid and taurocholic acid in follicular fluid, indicating bile acid metabolism changes affect both systemic circulation and the ovarian environment (15).

Elevated conjugated bile acids in PCOS may indicate hepatic metabolic changes, arising from increased conjugation or decreased deconjugation by the gut microbiota (16)(17). These acids are transported by hepatic transporters like OATP1B1 and OATP1B3, which prefer conjugated forms (18). Yoost et al. found taurocholic and taurodeoxycholic acids elevated in obese PCOS patients, highlighting a metabolic link (19). In this study, increased conjugated bile acids were seen across all PCOS patients, regardless of obesity, suggesting this is a core PCOS feature.

The significant elevation of cholic acid in the PCOS group highlights an important aspect of primary bile acid metabolism (15). Cholic acid is mainly produced via the classical (neutral) pathway, regulated by cholesterol 7 α-hydroxylase (CYP7A1) and sterol 12 α-hydroxylase (CYP8B1) (17). Higher cholic acid levels may indicate increased classical bile acid synthesis or changes in the CYP8B1 branch point that affects the cholic acid to chenodeoxycholic acid ratio. Notably, chenodeoxycholic acid levels did not differ significantly between groups, despite previous reports of elevated CDCA in PCOS patients (14). Yu et al. identified chenodeoxycholic acid as highly elevated in PCOS patients, with the highest VIP score in their OPLS-DA model (14). This discrepancy may reflect differences in study populations, metabolic phenotypes, or disease stage. The alternative pathway, producing mostly CDCA via CYP27A1, may be regulated differently across PCOS groups (17).

The absence of significant differences in secondary bile acids (deoxycholic acid and lithocholic acid) between groups suggests that intestinal microbial transformation of primary bile acids may not be substantially altered in the PCOS population studied (16)(17). Secondary bile acids are produced through 7 α-dehydroxylation of primary bile acids by specific gut bacteria, particularly Clostridium species (16)(20). The preservation of secondary bile acid levels contrasts with previous reports showing alterations in gut microbiota composition in PCOS patients, including changes in bacteria involved in bile acid metabolism (20).

This study’s functional bile acid indices reveal important insights into bile acid homeostasis in PCOS. The higher total conjugated bile acid pool in PCOS patients reflect increased taurocholic acid, glycocholic acid, and taurodeoxycholate (19). The conjugated-to-unconjugated ratio was also higher, indicating a shift toward conjugated forms due to enhanced hepatic conjugation, reduced intestinal deconjugation, or both (16). The secondary-to-primary bile acid ratio, reflecting intestinal microbial conversion, differed significantly between groups, suggesting possible changes in the gut microbiota (17). Qi et al. reported that women with PCOS have gut dysbiosis with more Bacteroides vulgatus and fewer bile acid metabolites like glycodeoxycholic and tauroursodeoxycholic acids (21). These changes may impact metabolic regulation and insulin sensitivity, as bile acids act as detergents and signaling molecules influencing glucose and lipid metabolism via nuclear and membrane receptors (22). Elevated conjugated bile acids may affect metabolic homeostasis through interactions with farnesoid X receptor (FXR) and the G protein-coupled bile acid receptor TGR5 (23).

Multivariate analysis strengthened the evidence of coordinated changes in bile acids associated with PCOS. Although unsupervised methods revealed some overlap among groups—attributable to shared bile acid pools—the overall profile shift indicates altered bile acid contributions. Deoxycholic acid emerged as a key factor distinguishing groups, even though it showed no significant difference in univariate tests, highlighting that bile acids function as part of an interconnected network. Recent PCOS metabolomics research supports these findings, showing that bile acids with minor individual differences can significantly impact multivariate discrimination (14).

Clinically, the correlation observed between bile acids and hormonal parameters suggest a relationship between bile acid metabolism and endocrine regulation in PCOS. The positive associations with cholic acid, secondary bile acids, and DHEA-S support a link to adrenal steroidogenesis, consistent with earlier research (14)(15). Additionally, correlations between secondary bile acids and the LH/FSH ratio suggest potential interactions between bile acid metabolism and hypothalamic–pituitary–ovarian axis dysregulation, though mechanistic studies are necessary to establish causality.

Identifying bile acid profiles in PCOS has significant clinical implications. It may serve as a diagnostic tool, especially when combined with markers such as testosterone. Yu et al. showed that a combination of chenodeoxycholic acid, lithocholic acid, and testosterone improves accuracy compared with testosterone alone (14). Bile acid measurements can also help stratify patients by metabolic risk, as suggested by their correlation with insulin resistance, lipids, and body composition, enabling targeted therapy and monitoring. The bile acid-gut microbiota axis is a promising therapeutic target; interventions such as probiotics, prebiotics, synbiotics, or faecal transplants may normalise metabolism and improve outcomes, as shown by Xue et al. in mice using inulin and metformin (24). Bile acid receptor agonists for FXR or TGR5 are under development for metabolic diseases and could be useful in PCOS (25). Zhang et al. highlight that lifestyle changes, especially adopting a Mediterranean diet and exercising, can positively influence the gut microbiota and bile acid metabolism and should be the first approach, with drugs for those not responding (17).

This is the first study in Indian ethnicity examining the relationship between bile acids and PCOS, offering valuable insights into this specific population. However, it has limitations: its cross-sectional design prevents causal inferences, and longitudinal research is needed for clearer understanding. Gut microbiota was not directly examined, so mechanistic links remain unclear; future studies should combine bile acid profiling with microbiome analysis. Dietary intake was not systematically assessed, though it influences bile acids and microbiota. The control group lacked glucose and insulin data, limiting comparisons of insulin resistance and metabolic parameters. Despite these limitations, the study provides a foundational step for future research and opens avenues for targeted interventions in this population.

This study reveals significant changes in serum bile acids in women with PCOS, indicating disrupted metabolism involving hepatic synthesis, conjugation, and gut microbiota. Key metabolites like deoxycholic acid, taurocholic acid, and cholic acid distinguish PCOS from healthy controls and correlate with steroidogenic parameters. These findings suggest that bile acid profiling, combined with microbiota analysis, could improve diagnosis, risk stratification, and targeted therapies, emphasizing the gut-liver-ovary axis as a promising treatment avenue.

## Abbreviations

PCOS: polycystic ovary syndrome
BMI: body mass index
LC-MS: liquid chromatography-mass spectrometry
LLOD: Lower limit of detection
LLOQ: Lower limit of quantification
PCA: Principal component analysis
OPLS-DA: Orthogonal partial least squares discriminant analysis
VIP: variable importance projection
FXR: farnesoid X receptor
TGR5: Takeda G-protein receptor 5
DCA: Deoxycholic acid
CDCA: Chenodeoxycholic acid
CA: Cholic acid
LCA: Lithocholic acid
GCA: Glycocholic acid
TCA: Taurocholic acid
TDCA: Taurodeoxycholic acid.

## Acknowledgements

We thank all the study participants and clinicians of Dr Nagori’s Institute, Ahmedabad and Gujarat University Health Center for their assistance. Furthermore, we are deeply grateful to the Scheme of Developing HighQuality Research (SHODH) Department of Education, Government of Gujarat, India, for providing fellowship to Jalpa Patel (SHODH Reference Id: 202001380093) and CSIR-UGC-NET, providing companionship to Hiral Chaudhary (Ref No: CSIR UGC NET 947).

## Author contributions

JP assisted with data collection, writing, and manuscript preparation. HC and SP assisted with data collection. BP and RJ drafted and reviewed this manuscript. All authors approved the submitted version, take personal responsibility for their contributions, and commit to properly investigating any questions about the accuracy or integrity of the work, with findings documented in the literature.

## Funding

This research was funded by the Knowledge Consortium of Gujarat (KCG) (grant number KCG/0062/10/2025/ RFS457)

## Data availability

The datasets used and analyzed during the current study are available from the corresponding author upon reasonable request.

## Declarations

### Ethics approval and consent to participate

The Institutional Ethics Committee (IEC) (GU-IEC(NIV)/02/Ph.D./007) of the University School of Sciences, Gujarat University, approved this study. All participants gave informed consent. All research work was performed in accordance with relevant guidelines and regulations. These guidelines and considerations align with the ethical approval requirements of the Helsinki Declaration, which emphasizes the protection of human subjects in research.

### Consent for publication

Not applicable.

### Competing interests

The authors declare no competing interests.

## Notes

### Competing Interest Statement

The authors have declared no competing interest.

