## Supplemental Figure 1 for "Serum Bile Acid Dysregulation in Polycystic Ovary Syndrome: Quantitative Insights from Mass Spectrometry-Based Profiling"

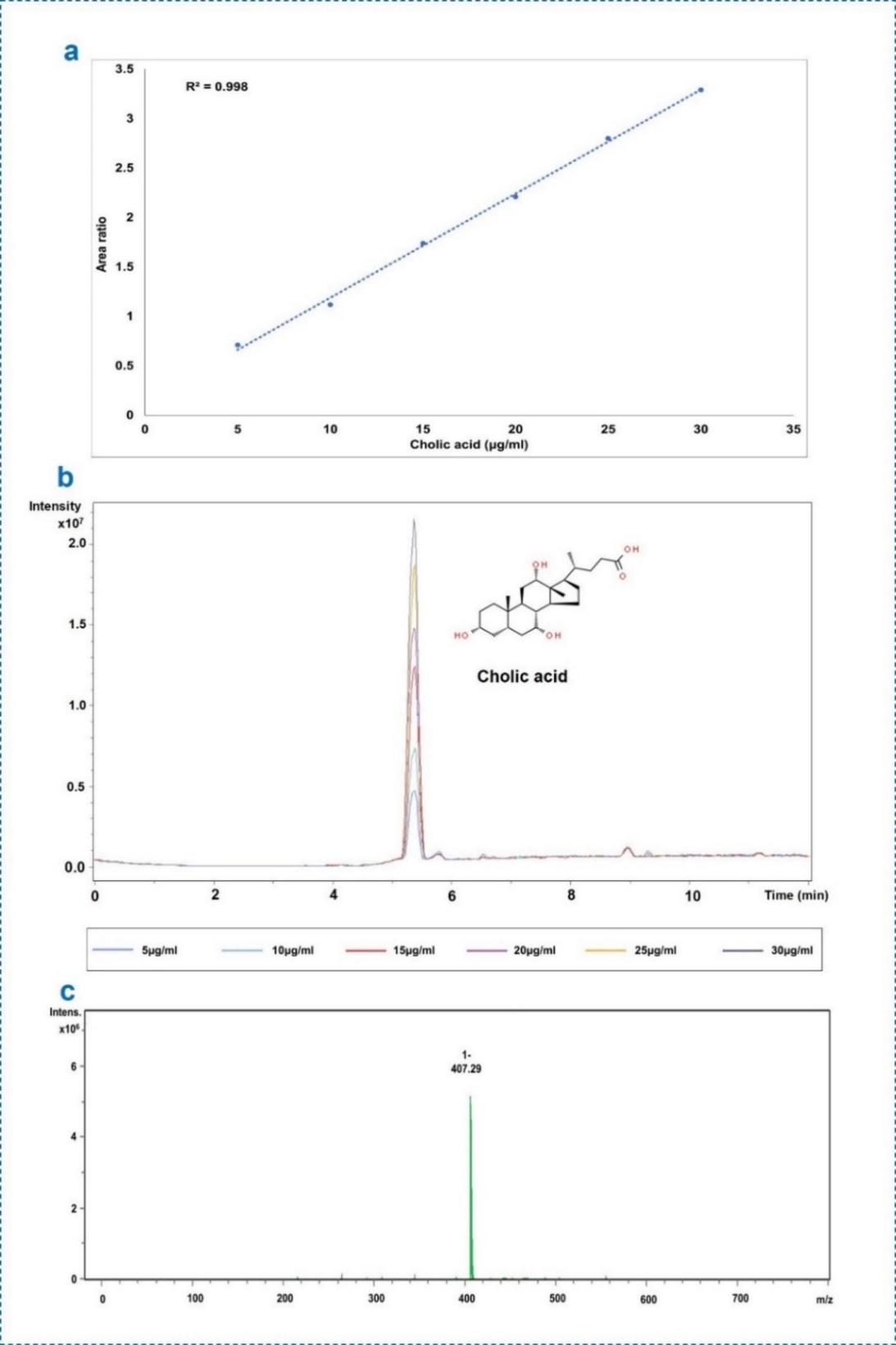

**Supplementary Figure 1.** Chromatographic quantification of Cholic Acid. (a) Calibration curve illustrating the linear relationship between Cholic acid concentration and area ratio (R² = 0.998). (b) Chromatogram showing distinct peaks at a retention time of approximately 5.4 minutes for different concentrations of Cholic acid. (c) Mass spectrum confirming the molecular identity of Cholic acid with a peak at m/z 407.29.
