## Supplemental figure 2 for "Serum Bile Acid Dysregulation in Polycystic Ovary Syndrome: Quantitative Insights from Mass Spectrometry-Based Profiling"

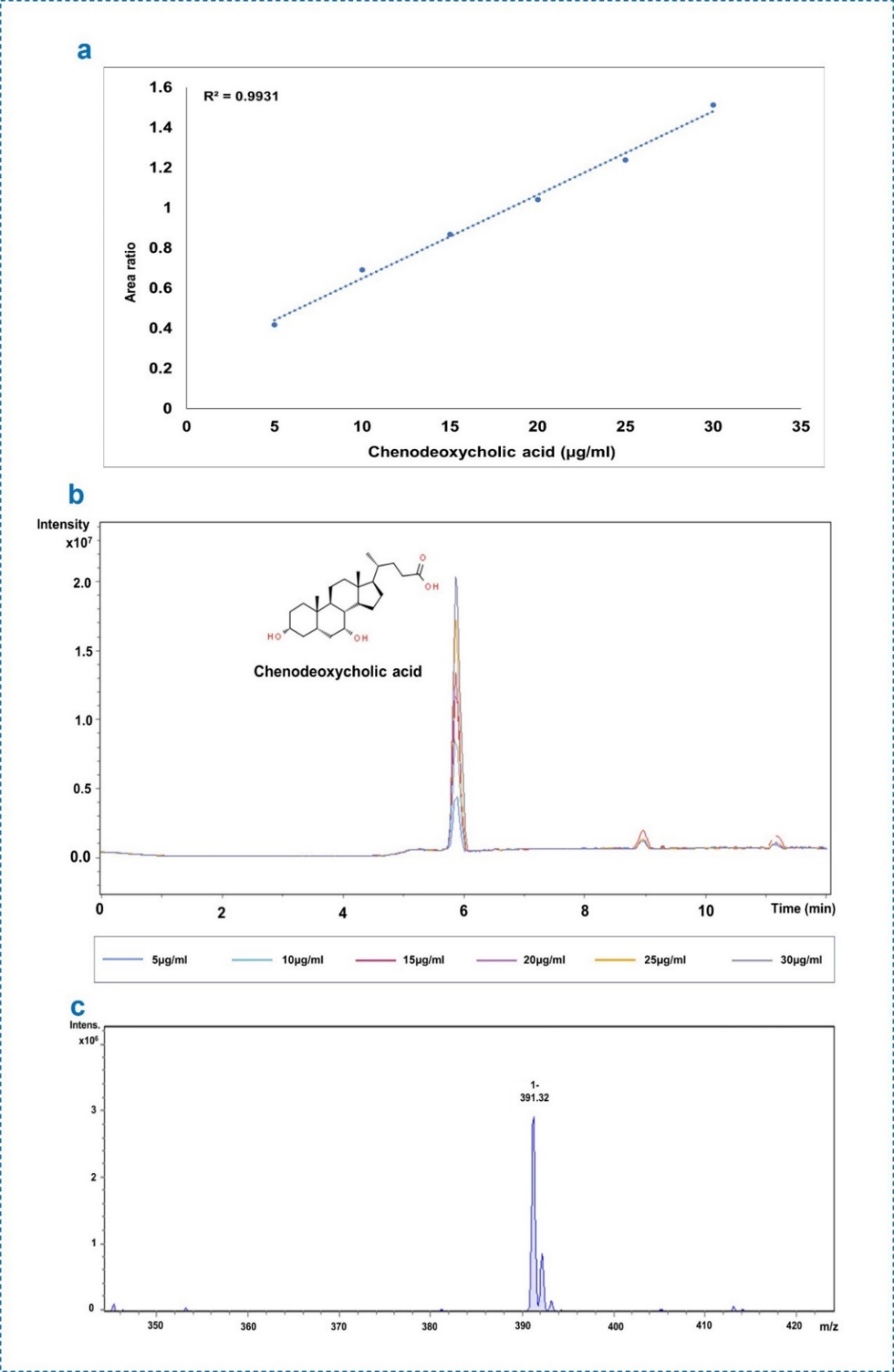

**Supplementary Figure 2.** Chromatographic representation of quantification of Chenodeoxycholic Acid. (a) Calibration curve demonstrating the linearity of Chenodeoxycholic acid concentration and area ratio (R² = 0.9931). (b) Chromatogram showing consistent peaks at a retention time of 5.9 minutes for varying concentrations. (c) Mass spectrum identifying Chenodeoxycholic acid with a peak at m/z 391.32.
