## Supplemental figure 3 for "Serum Bile Acid Dysregulation in Polycystic Ovary Syndrome: Quantitative Insights from Mass Spectrometry-Based Profiling"

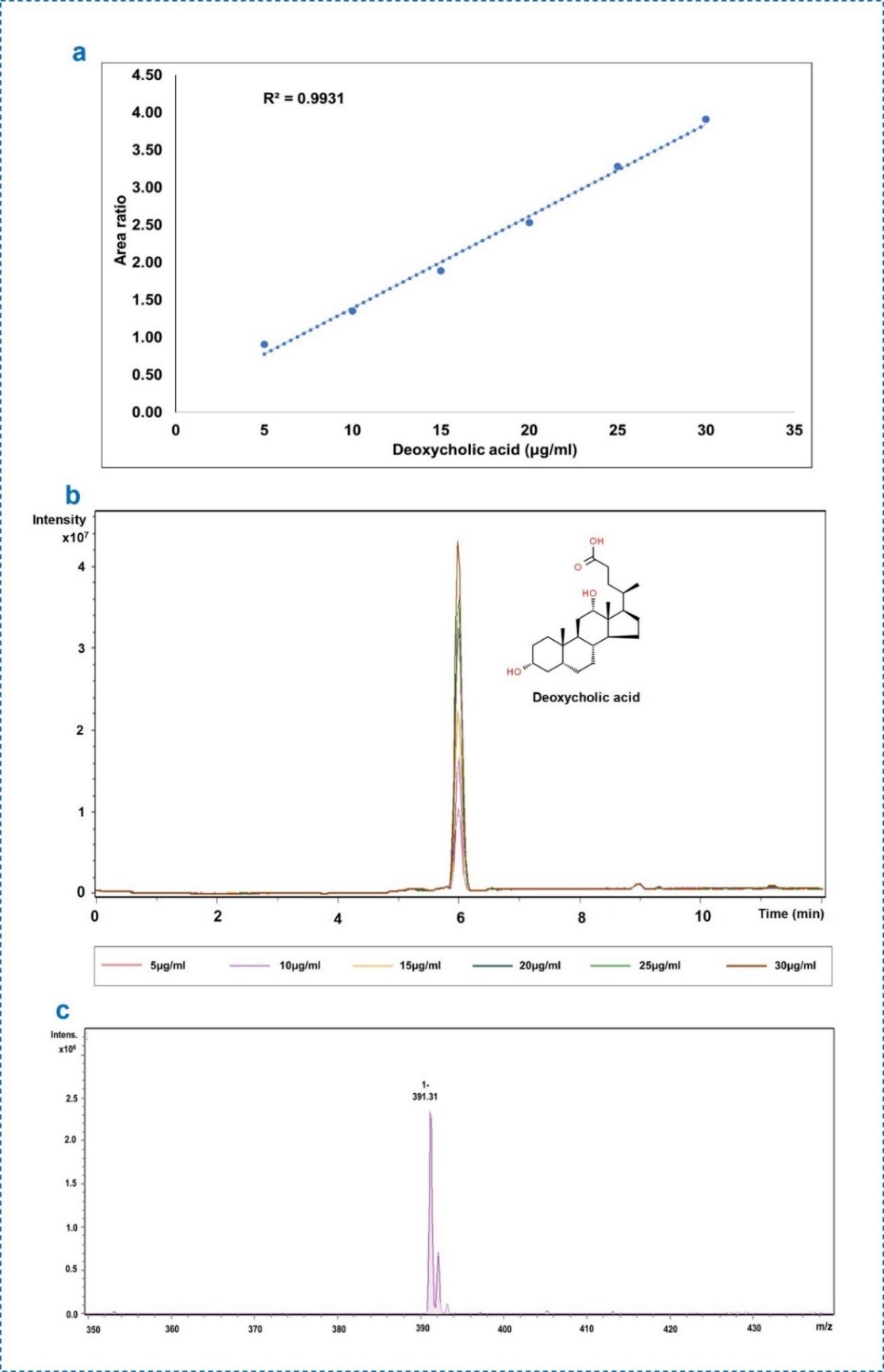

**Supplementary Figure 3.** Chromatographic quantification of Deoxycholic Acid. (a) Calibration curve showing a linear relationship between Deoxycholic Acid concentration and area ratio (R² = 0.9931). (b) Chromatogram with distinct peaks at approximately 6 minutes. (c) Mass spectrum confirming the identity of Deoxycholic Acid with a peak at m/z 391.31.
