## Supplemental figure 4 for "Serum Bile Acid Dysregulation in Polycystic Ovary Syndrome: Quantitative Insights from Mass Spectrometry-Based Profiling"

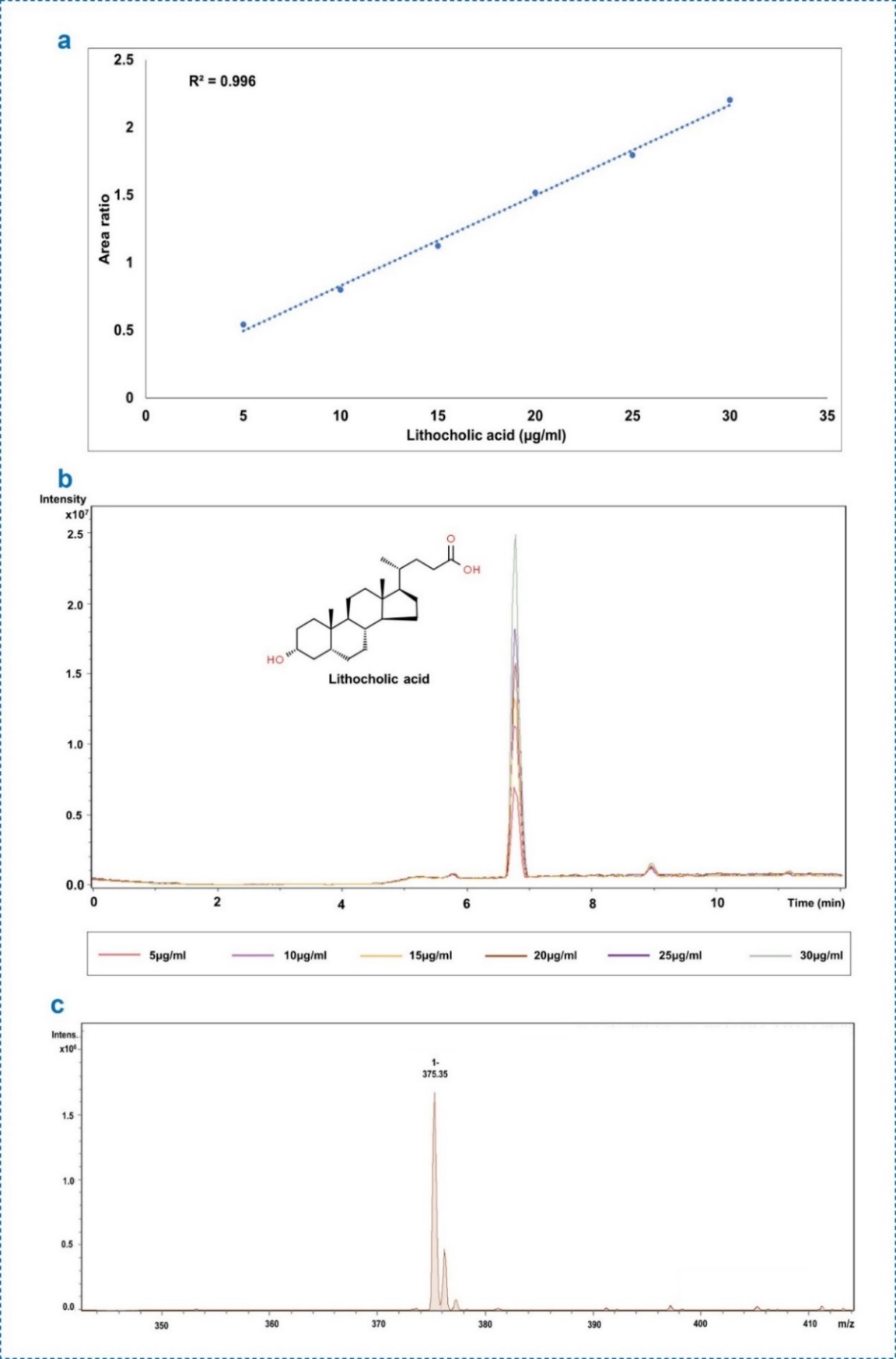

**Supplementary Figure 4.** Chromatographic representation of Quantification of Lithocholic Acid. (a) Calibration curve showing high linearity for Lithocholic acid concentration and area ratio (R² = 0.996). (b) Chromatogram displaying consistent peaks at 6.8 minutes. (c) Mass spectrum identifying Lithocholic acid with a peak at m/z 375.35.
