## Supplemental figure 5 for "Serum Bile Acid Dysregulation in Polycystic Ovary Syndrome: Quantitative Insights from Mass Spectrometry-Based Profiling"

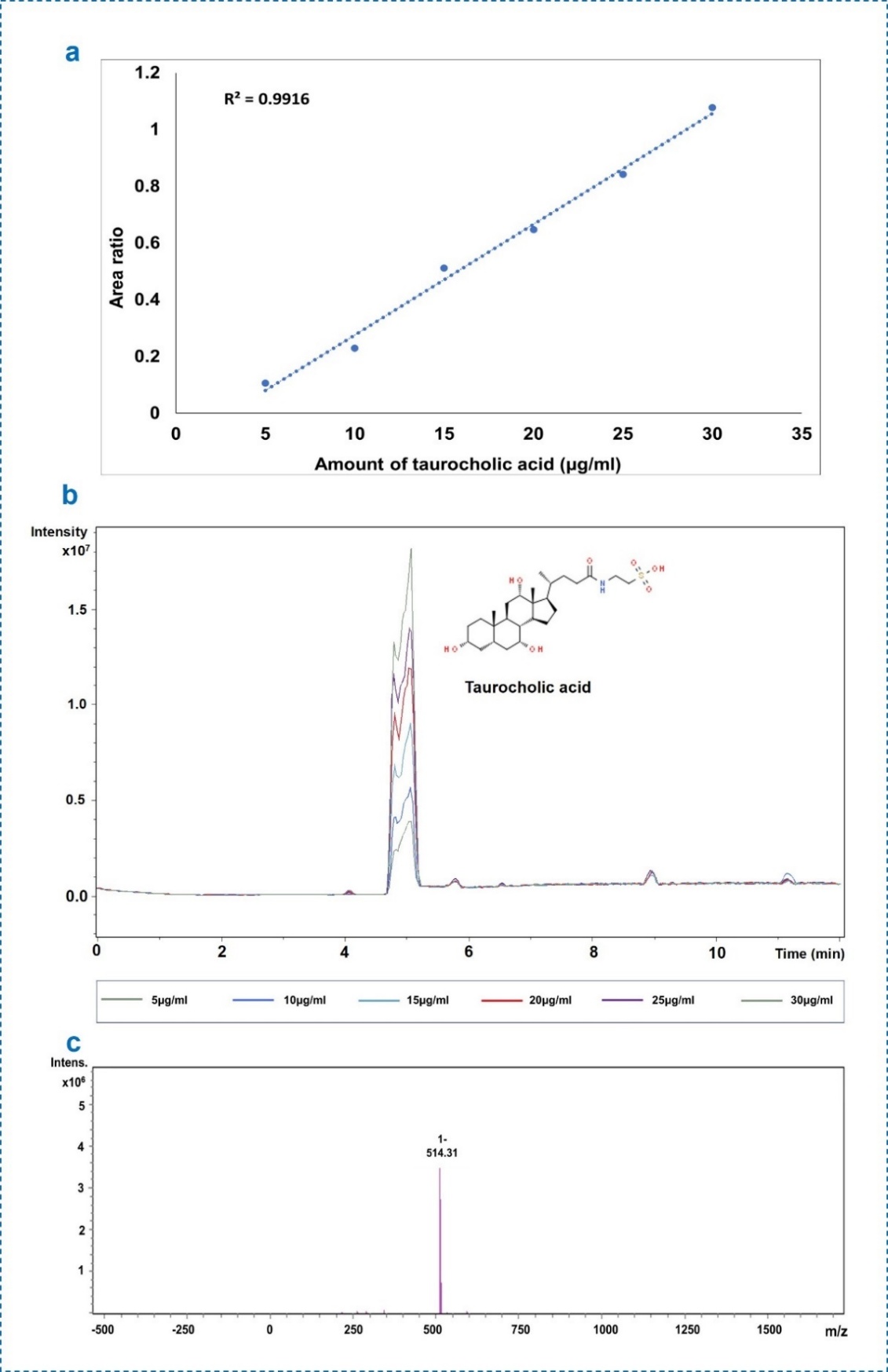
**Supplementary Figure 5.** Chromatographic quantification of Taurocholic Acid. (a) Calibration curve showing the linear relationship between Taurocholic Acid concentration and area ratio (R² = 0.9916). (b) Chromatogram with distinct peaks at approximately 4.8 minutes. (c) Mass spectrum confirming the identity of Taurocholic Acid with a peak at m/z 514.31.
