## Supplemental figure 6 for "Serum Bile Acid Dysregulation in Polycystic Ovary Syndrome: Quantitative Insights from Mass Spectrometry-Based Profiling"

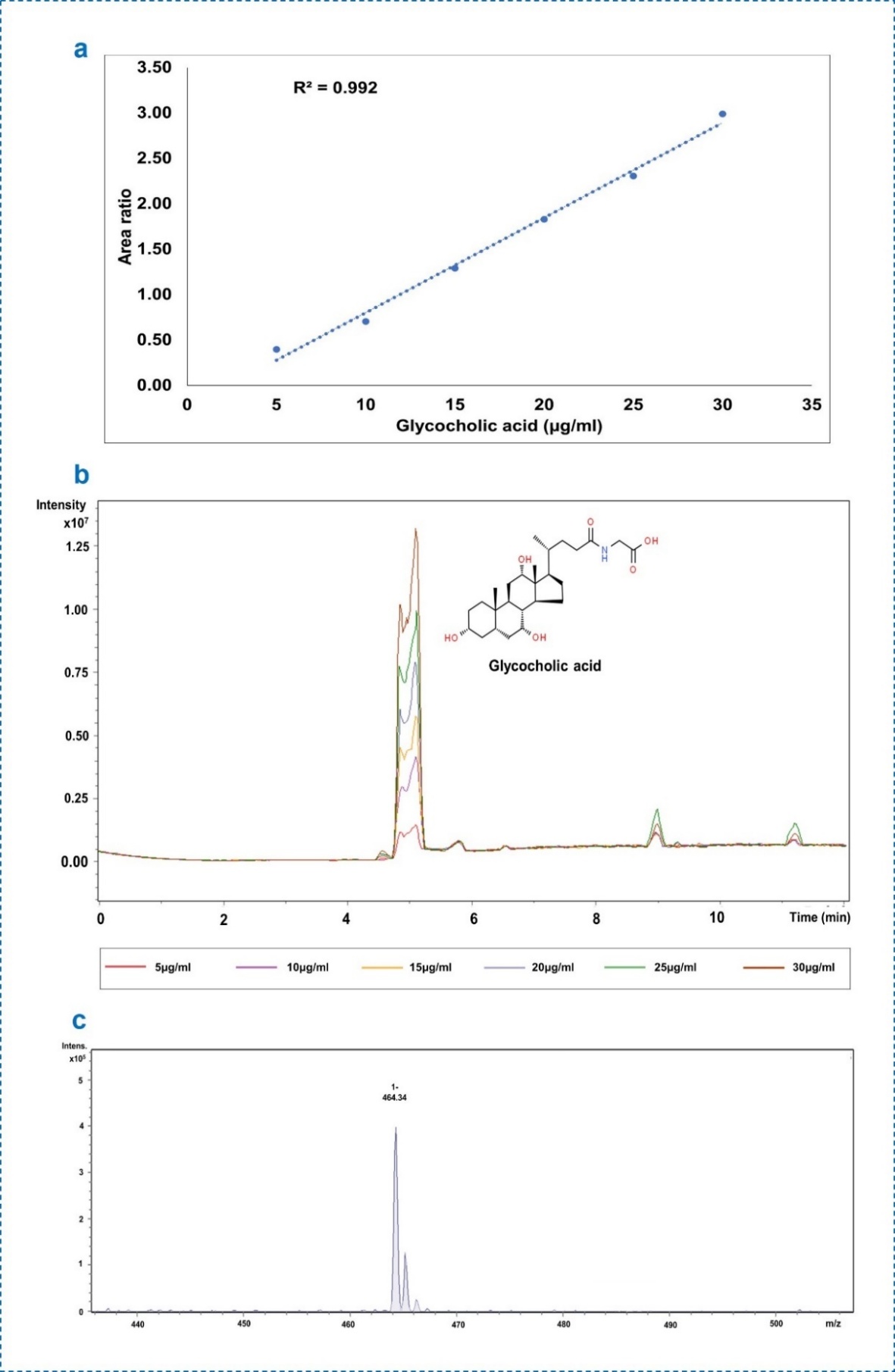
 **Supplementary Figure 6.** Chromatographic representation of quantification of Glycocholic Acid. (a) Calibration curve showing high linearity between Glycocholic Acid concentration and area ratio (R² = 0.992). (b) Chromatogram showing distinct peaks around 4.9 minutes. (c) Mass spectrum identifying Glycocholic Acid with a peak at m/z 464.34.
