## Supplemental figure 7 for "Serum Bile Acid Dysregulation in Polycystic Ovary Syndrome: Quantitative Insights from Mass Spectrometry-Based Profiling"

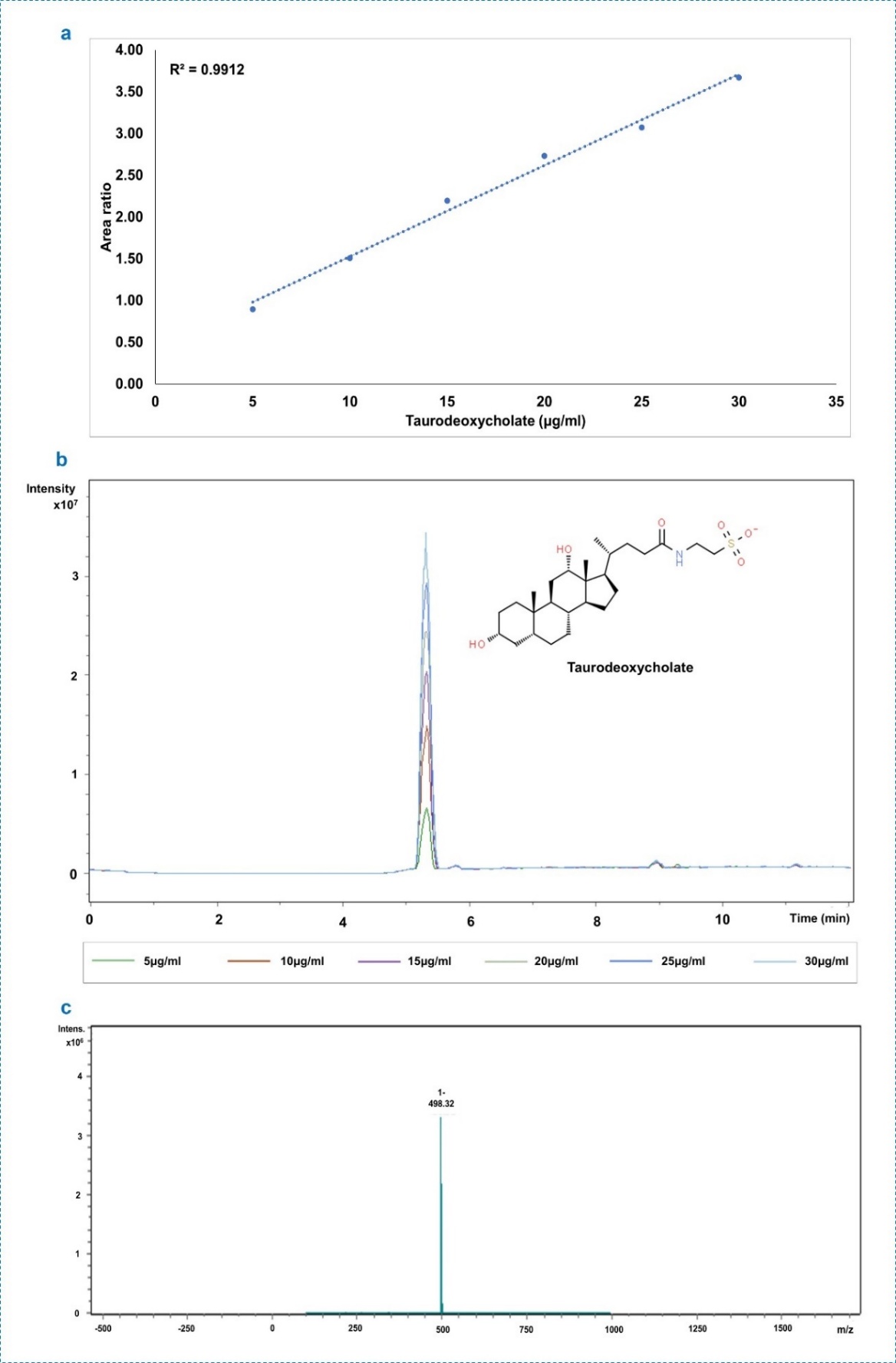
**Supplementary Figure 7.** Chromatographic representation of quantification of Taurodeoxycholate. (a) Calibration curve demonstrating a linear relationship for Taurodeoxycholate concentration and area ratio (R² = 0.9912). (b) Chromatogram with consistent peaks at approximately 5.3 minutes. (c) Mass spectrum confirming the identity of Taurodeoxycholate with a peak at m/z 498.32.
